# IL-6 blockade suppresses the blood-brain barrier disorder, leading to prevention of onset of NMOSD

**DOI:** 10.1101/2021.01.28.428564

**Authors:** Yukio Takeshita, Susumu Fujikawa, Kenichi Serizawa, Miwako Fujisawa, Kinya Matsuo, Joe Nemoto, Fumitaka Shimizu, Yasuteru Sano, Haruna Tomizawa-Shinohara, Shota Miyake, Richard M. Ransohoff, Takashi Kanda

## Abstract

Neuromyelitis optica spectrum disorder (NMOSD) is an autoimmune astrocytopathy caused by antibodies against the aquaporin 4(AQP4) in end-feet of astrocytes. Breakdown of the blood–brain barrier (BBB) allowing ingress of AQP4 antibodies into the central nervous system (CNS) plays a key role in NMOSD. Although IL-6 blockade therapies such as satralizumab are effective in NMOSD, the therapeutic mechanism of IL-6 blockade, especially with respect to BBB disruption, are not fully understood because of the lack of the human models that are specialized to evaluate the BBB function.

We constructed new in vitro human BBB models for evaluating continued barrier function, leukocyte transmigration and intracerebral transferability of IgGs utilizing the newly established triple co-culture system. In vitro and vivo experiments revealed that NMO-IgG increased intracerebral transferability of satralizumab, and that satralizumab suppressed the NMO-IgG-induced transmigration of T cells and barrier dysfunction. These results suggest that satralizumab, which can pass through the BBB in the presence of NMO-IgG, suppresses the barrier dysfunction and the disrupting controlled cellular infiltration at the BBB, leading to prevention of onset of NMOSD.

**One sentence summary:** Satralizumab and IL-6 blockade prevent lymphocyte migration and barrier dysfunction induced by NMO-IgG in EAE and novel triple co-culture BBB models.

## Introduction

Neuromyelitis optica spectrum disorder (NMOSD) is an inflammatory disease of the central nervous system (CNS) associated with recurrent optic neuritis and longitudinally extensive transverse myelitis (*1, 2*). A major characteristic of NMOSD is the development of antibodies against the water channel aquaporin-4 (AQP4), which is expressed mainly in astrocytic foot processes (*3, 4*). The IgG plasma fraction of NMOSD patients (NMO-IgG) contains anti-AQP4 antibodies, leading to complement- and antibody-dependent cellular cytotoxicity of astrocytes (*5-7*). In order for peripherally produced anti-AQP4 antibodies to gain access to their target in the CNS, they need to penetrate the blood–brain barrier (BBB) (*8*). Accordingly, disruption of the BBB—which allows influx of humoral factors including autoantibodies through the dysfunctional barrier and infiltration of inflammatory cells by disruption of controlled cellular infiltration—is considered to be the first and key step in the pathogenesis of NMOSD (*9-11*). In fact, the ratio of cerebrospinal fluid (CSF) to serum albumin (CSF:serum albumin ratio), which is a marker of BBB permeability, is correlated with the clinical severity of NMOSD (*12, 13*).

Levels of interleukin-6 (IL-6) are elevated in the CSF and serum of patients with NMOSD as compared with levels in patients with multiple sclerosis or non-inflammatory neurologic disease (*14-16*). In earlier studies (*11, 17*), we found that anti-GRP78 and anti-AQP4 autoantibodies in NMO-IgG were important factors in the breakdown of the BBB via induction of IL-6 expression in astrocytes. In recent *ex-vivo* experiments, inhibition of IL-6 signaling was shown to inactivate the effector functions of plasmablasts, which are a major source of NMO-IgG in the peripheral blood (*18*). In terms of BBB disruption, IL-6 dose- and time-dependently decreases the expression of endothelial tight junction proteins and increases permeability in human brain microvascular endothelial cells (*19*). In patients with NMOSD, increased IL-6 in the CSF is positively correlated with the CSF:serum albumin ratio (*20*). Overall, these reports suggest that IL-6 signaling pathways are involved in the pathogenesis of NMOSD. In fact, clinical research has demonstrated that treatment with anti-IL-6 receptor antibody ameliorates the disease in patients with NMOSD (*21, 22*).

Satralizumab is a humanized immunoglobulin G subclass 2 (IgG2) monoclonal antibody against IL-6 receptors; it specifically binds to both membrane-bound and soluble forms of IL-6 receptors and blockades IL-6 signaling pathways (*23*). The constant and variable regions of satralizumab are engineered to have pH-dependent binding to IL-6 receptors, increased affinity to the neonatal Fc receptor, and a lower isoelectric point to extend the elimination half-life of the drug in plasma (*24, 25*). Importantly, it is reported that satralizumab showed beneficial effects such as a lower risk of relapse than with placebo among patients with NMOSD in an international, randomized, double-blind, placebo-controlled, phase 3 trial (*26*). However, the therapeutic mechanisms of satralizumab, especially those with respect to BBB disruption, still remain unknown because there are no ideal *in vivo* or *in vitro* BBB models with which to explore NMOSD pathogenesis.

To address this pertinent issue, we propose some important properties that will be necessary for an *in vitro* BBB model that is more robust than those currently available (*27*). First, the model should include human cells that maintain both physiological and morphological BBB properties *in vitro*. Second, endothelial cells should be co-cultured with other BBB cells such as astrocytes and pericytes. Third, the model should allow the transendothelial migration of inflammatory cells under physiologically relevant shear forces. Additionally, the model should allow recovery of transmigrated cells for further analysis of BBB function, including real-time monitoring of transendothelial electrical resistance (TEER) and the measurement of microvolumes of humoral factors or IgG translocation across the BBB.

In order to construct *in vitro* BBB models with these properties, we utilized human brain microvascular endothelial cells (hECs; TY10), human astrocytes (hASTs) with AQP4 expression, and human pericytes (hPCTs), each of which was conditionally immortalized by transfection with temperature-sensitive SV40 (simian virus 40) large T antigen (ts-SV40-LT) and which retain both their physiological and morphological BBB properties (*28, 29*). We earlier constructed a flow-based *ex-vivo* models of hECs co-cultured with hASTs that enables us to evaluate the transmigration of leukocytes across the endothelium under shear forces (*11, 30, 31*). However there were no ideal triple co-cultured *in vitro and ex-vivo* BBB model in which pericytes and the end-feet of astrocytes can directly contact endothelial cells. Then it was impossible to evaluate the barrier function, leukocyte transmigration and intracerebral transferability to reveal NMOSD pathogenesis

In the present study, we constructed functional *in vitro* static and *in ex-vivo* flow-based models utilizing the newly established triple co-culture system of temperature-sensitive conditionally immortalized human BBB cell lines (endothelial cells [hECs], pericytes [hPCTs], and astrocytes with AQP4 expression [hASTs]) in order to explore the effects of NMO-IgG on the BBB. The new static *in vitro* model allowed long-term measurement of real-time TEER by means of an automated cell monitoring system and measurement of microvolumes of IgG translocation through the BBB. The new *in ex-vivo* flow-based model enabled us to evaluate leukocyte transmigration across the BBB. Then, using these structured *in vitro* BBB models, we evaluated the effects of satralizumab on BBB disruption caused by NMO-IgG. In addition to the *in vitro* assays, we also assessed the effects of an anti-IL-6 receptor antibody for mice (MR16-1) on *in vivo* BBB disruption in mice with experimental autoimmune encephalomyelitis (EAE). These mice are used as an animal model of CNS autoimmune diseases in which IL-6 concentration in the spinal cord dramatically increases (*32*).

## Results

### Construction newly in vitro BBB models with triple co-culture system of temperature-sensitive conditionally immortalized human BBB cell lines

We utilized human brain microvascular endothelial cells (hECs; TY10), human astrocytes (hASTs) with AQP4 expression, and human pericytes (hPCTs), each of which was conditionally immortalized by transfection with temperature-sensitive SV40 (simian virus 40) large T antigen (ts-SV40-LT) and which retain both their physiological and morphological BBB properties(Fig. 1A). In transfected these cell lines, cells are driven to continue proliferating at 33°C because activated Ts-SV40 LT inhibited P53 and Rb. On the other hand, cells could differentiate into mature cells at 37 °C because Ts-SV40 LT is inactivated and exhibit growth arrest (Fig.1 B). To construct triple co-culture systems of these temperature-sensitive conditionally immortalized human BBB cell lines, multiple steps were organized by including cell-culture on Upcell dish with coated temperature-responsive polymer (Fig. C and D). Firstly, hASTs were cultured on abluminal side of insert membranes having 3 μm pores and incubated for 24 hours so that some astrocytic end-feet with AQP4 could protrude through the membrane pores (Fig. 1 E and F). Secondly, after culturing of hPCTs on the luminal side of membranes (Fig. 1 G), hASTs and hPCTs were co-cultured at 33 °C for 24 hours. Thirdly, hECs were cultured on Upcell dish with coated temperature-responsive polymer, which can achieve sheet-like detachment of confluent cells and extra-cellular matrix by temperature-shifting to 20°C (Fig. 1 I and H). Then sheet-like detachment of confluent hEC were transferred onto the hPCTs. After co-cultuting of these cell lines at 33 °C for 24 hours, they differentiated into mature cells under the condition of 37 °C. Confocal 3D analysis with living staining of each cell line showed that multi-cultured insert constituted the five-layer structures which is consisted of hEC, hPCT, astrocytic endfeet, membrane, and hAST(Fig.1 J 1-5). Some end-feet protruded through the membrane pores. hPCTs and end-feet of hASTs were close to the hEC layer.

**Figure 1.**
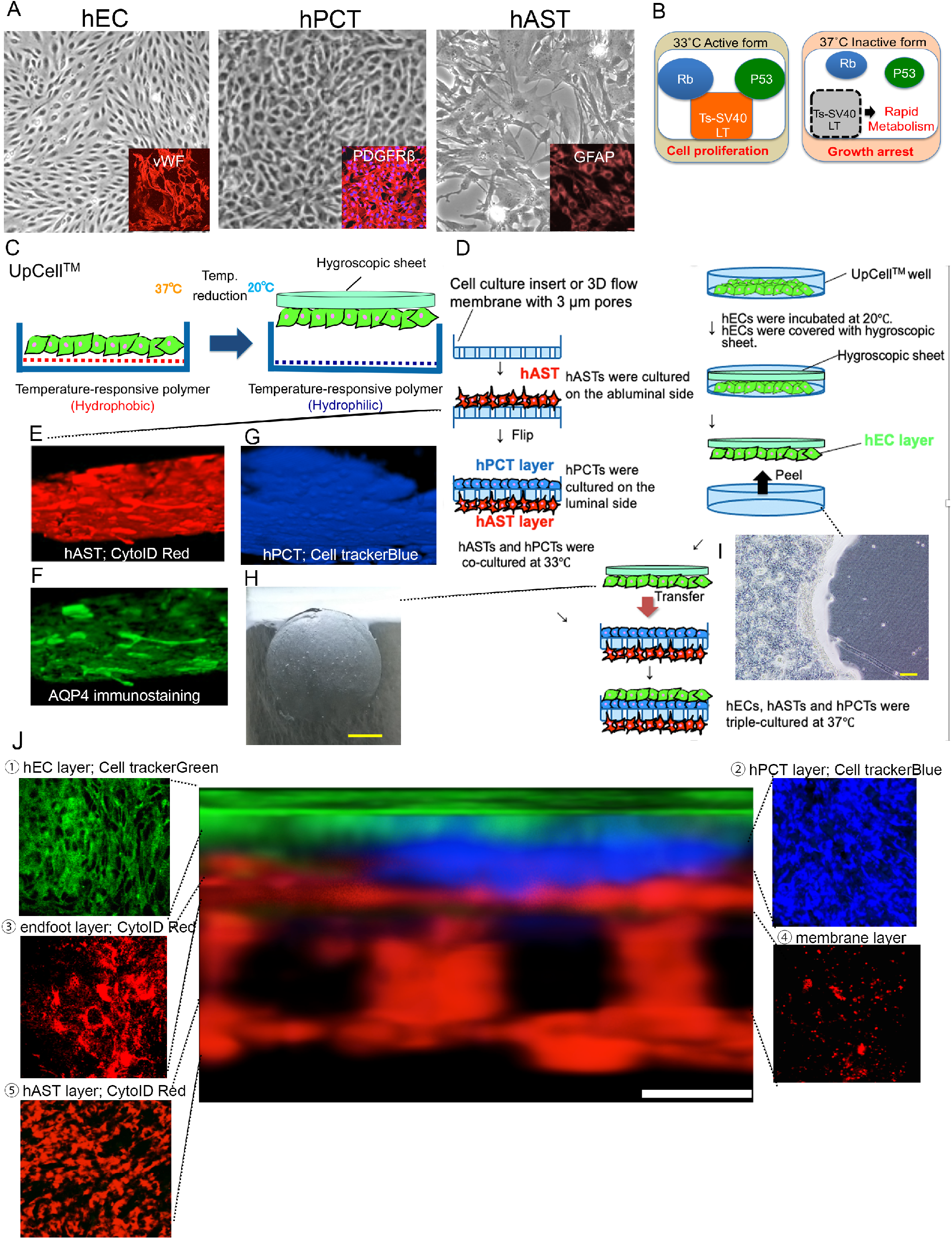
Construction newly *in vitro* BBB model with triple co-culture system of temperature-sensitive conditionally immortalized human BBB cell lines. A) Establishment of the BBB cell lines that maintains the BBB properties. Morphology of hEC is spindle-shape. hEC expressed vWF as lineage marker of endothelium. Morphology of hPCT is cobblestone-shape. hPCT expressed PDGFβ as lineage marker of pericyte. Morphology of hAST is star-shape. hAST expressed GFAP as lineage marker of hAST. B) Three conditionally immortalized human cell lines were transfected with temperature sensitive SV40 large T antigen (Ts-SV40 LT). At 33°C, actived Ts-SV40 LT binds and inhibits p53 and Rb, which are strong tumor suppressors, leading to continuous cell proliferation. At 37 °C, inactivated Ts-SV40 LT exhibit growth arrest, leading to differentiate into mature cells. C) The UpCell™ technology. Temperature-responsive polymer is immobilized on the surface of UpCell™ well. The polymer grafted surface shows reversible hydrophobic - hydrophilic property across the threshold temperature of 32°C, cell-sheet is detached from the dish without harmful enzymes (e.g. trypsin or dispase) and attached the hygroscopic sheet to transfer. D) Multistep of triple co-culture system. hASTs were cultured on the abluminal side of cell culture insert or 3D flow membrane with 3μm pores. After flipping of cultured insert, hPCTs were cultured on the luminal side. hAST and hPCTs were co-cultured at 33°C. hEC were cultured on UpCell dish. After incubation at 20°C, The sheet of confluent hECs was detached and transferred onto the hPCTs co-cultured with hASTs on the insert. E,F) Astrocytic endfoot with AQP4 on cultured membrane. living-stained hAST cultured on membrane. Confocal 3D analysis from luminal side of membrane with living staining of hAST(E) and immunostaining of AQP4(F) showed some astrocytic end-feet protruded through the membrane pores. G) hPCT were co-cultured on the luminal side of the membrane. Confocal 3D analysis from luminal side of membrane with living staining of hPCT showed hPCT layer was on the membrane without any tentacles like astrocytic end-feet. H-I) Sheet-like detachment (H) and traces of peeling on dish of confluent hECs (I). H, bar = 200 μm. I, ber= 10 mm. J) Confocal 3D analysis with living staining of each cell lines. Multi-cultured insert constituted the five-layer structures which is consisted of hEC, hPCT, astrocytic endfeet, membrane, and hAST (1-5). Some end-feet protruded through the membrane pores. hPCTs and end-feet of hASTs were close to the hEC layer. bar = 5 μm

### Construction of ex vivo BBB model with triple co-culture system for leukocyte transmigration and effect of satralizumab on NMO-IgG-induced transmigration of leukocytes in ex-vivo

To evaluate the effect of satralizumab on NMO-IgG-induced transmigration of leukocytes, we constructed a flow-based dynamic BBB model incorporating hEC/hPCT/hAST triple co-culture, which allows further investigation of leukocyte transmigration under flow (Fig.2 A-D). After exposing the endothelial cell side (vascular side) and the astrocyte side (brain parenchymal side) to satralizumab plus NMO-IgG or to NMO-IgG alone (Fig.2. E), we counted the total numbers of all migrating cells and the numbers of phenotyped cells and compared their migrations relative to those with Control IgG. Application of NMO-IgG increased the migrations of all peripheral blood mononuclear cells (PBMCs) and CD4^+^, CD8^+^, and CD19^+^ cells relative to their migrations with Control IgG, and the application of NMO-IgG plus satralizumab significantly suppressed this increase in the relative numbers of migrating PBMCs and CD4^+^ and CD8^+^ cells (Fig. 2 F).

**Figure 2.**
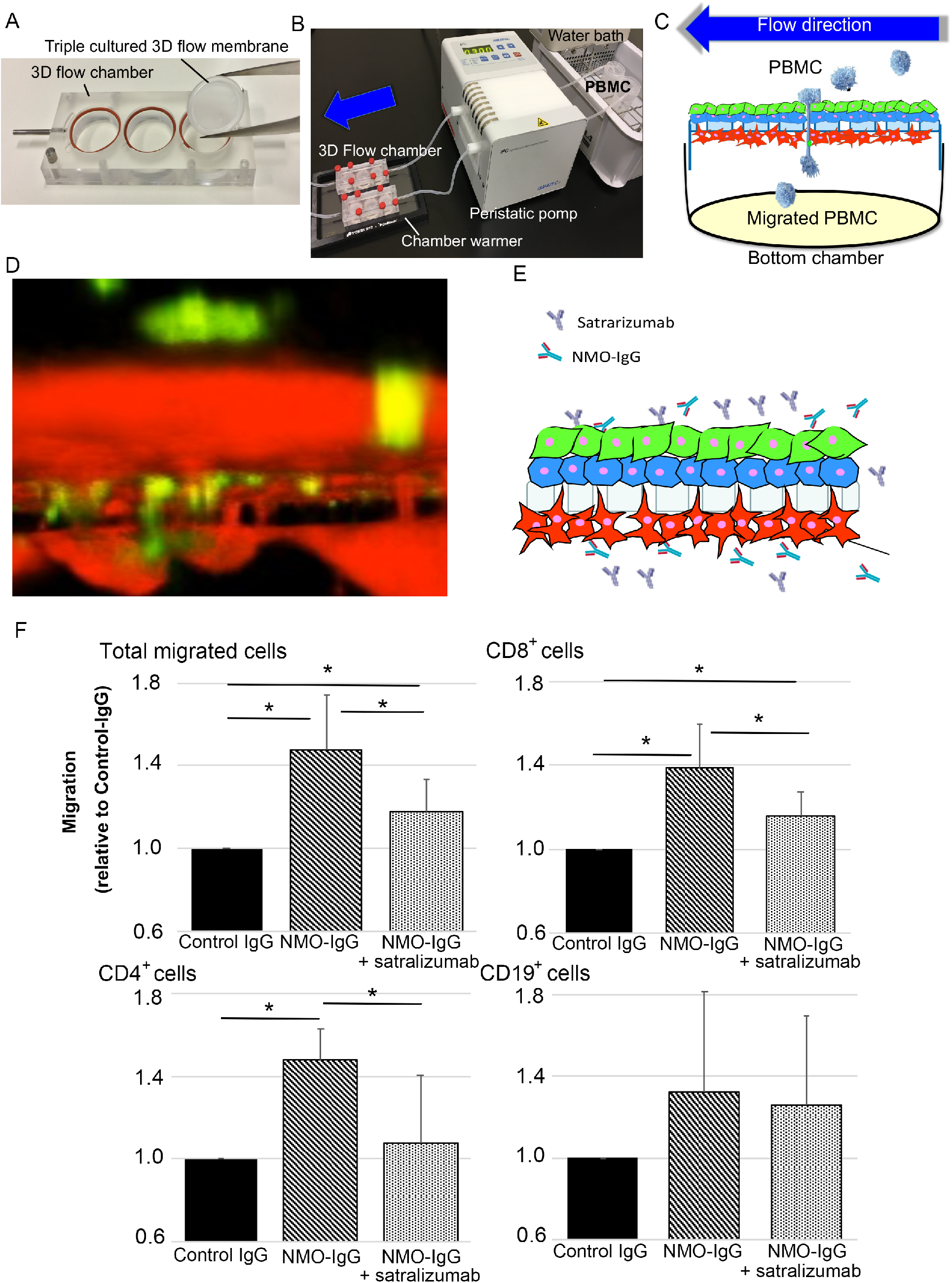
Construction of *ex-vivo* BBB model with triple co-culture system for leukocyte transmigration and effect of satralizumab on NMO-IgG-induced transmigration of leukocytes *in ex-vivo*. A) 3D flow chamber and 3D flow membrane. the triple-cultured membranes were transferred in 3D chamber. B) Whole set up of migration assay under shear forces. Normal human peripheral blood mononuclear cells (PBMC) flowed onto luminal side with physiological shear force by peristatic pomp. C) Schema of 3D flow chamber in transmigration assay. Total migrated cells were recovered from the bottom chamber and enumerated. D) Confocal 3D analysis with living staining of BBB cell lines and PBMCs in transmigration assay. PBMC were stained with Cell trackerGreen and all human BBB cell lines (hECs, hPCTs, and hASTs) were stained with CytoID Red. Some PBMCs protruded through the membrane pores. E) Schema of leukocyte transmigration assay with NMO-IgG and/or Satralizumab. F) Flow-based leukocyte transmigration assays utilizing a 3D flow Chamber showed that application of NMO-IgG increased the numbers of total migrating PBMCs and CD4^+^, CD8^+^, and CD19^+^ cells relative to numbers migrating with Control IgG, and that the application of NMO-IgG plus satralizumab significantly suppressed that increase. *\*P* < 0.05 by unpaired t-test (*n* = 6 per group). All data are expressed as mean and SEM.

### Effects of IL-6 receptor blockade on clinical signs and on transmigration of leukocytes in EAE mice

Administration of MR16-1 (anti-IL-6 receptor antibody for mice) on Day 7 after induction of experimental autoimmune encephalomyelitis significantly and strongly prevented the onset of clinical signs in EAE mice (Fig. 3A). We then confirmed the effect of MR16-1 on leukocyte transmigration in EAE mice. On Day 15, the number of CD4^+^ T cells markedly increased in the spinal cords of EAE mice, whereas these cells were almost undetected in the spinal cords of Control mice (Control, 33.7 ± 21.5 [mean ± SEM] cells/spinal cord slice; EAE, 628.3 ± 196.5 cells/spinal cord slice) (Fig. 3B). Administration of MR16-1 on Day 7 significantly prevented the migration of CD4^+^ T cells into the spinal cord (EAE + MR16-1, 40.3 ± 17.7 cells/spinal cord slice) (Fig. 3B). To exclude the possibility that the impact of MR16-1 on clinical signs and on leukocyte migration into the spinal cord was secondary to changes in the immune response, we evaluated the effect of MR16-1 on T cell differentiation in EAE mice. In splenocytes, Th1 cells and FoxP3-positive regulatory T cells were significantly increased on Day 16 in EAE mice (Fig. 3, C and E). Th17 cells showed a tendency, but not significantly, to increase in EAE mice (Fig. 3D). Administration of anti-IL-6 receptor antibody on Day 7 did not affect the induction of these in EAE mice (Fig. 3, C to E).

**Figure 3.**
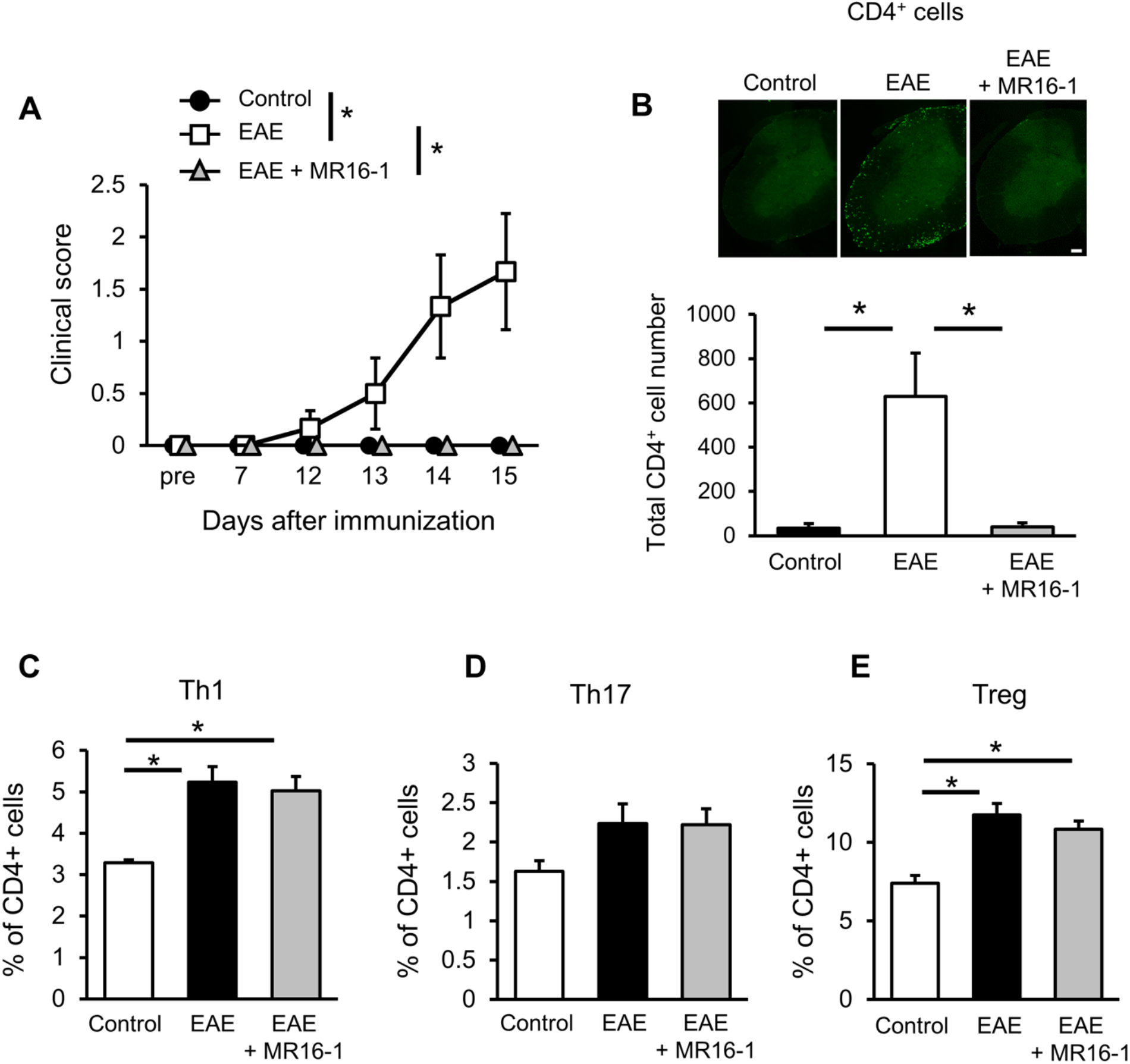
Effects of IL-6 receptor blockade on lymphocyte migration into the spinal cord *in vivo*. (**A**) Anti-IL-6 receptor antibody (MR16-1) was administered on Day 7 after immunization. Anti-IL-6 receptor antibody significantly prevented the onset of clinical signs in EAE mice. *\*P* < 0.05 by two-way ANOVA (*n* = 3-6 per group). (**B**) Anti-IL-6 receptor antibody suppressed lymphocyte migration into the spinal cords of EAE mice. Representative images showing immunohistochemical staining for CD4^+^ cells in the spinal cord on Day 15 after immunization. The number of CD4^+^ T cells was markedly increased in the spinal cord of EAE mice. Anti-IL-6 receptor antibody administered on Day 7 after immunization significantly prevented this increase. \**P* < 0.05 by Tukey’s multiple comparison test (*n* = 3-6 per group). Scale bar = 100 μm. (**C**) The induction of Th1 cells and (**E**) FoxP3-positive regulatory T cells was significantly upregulated on Day 16 after immunization. There was a tendency, but not significantly, for Th17 cells to increase in EAE mice (**D**). Administration of anti-IL-6 receptor antibody on Day 7 after immunization did not change the induction of these in EAE mice. * *P* < 0.05 by Tukey’s multiple comparison test (*n* = 4-8 per group). All data are expressed as mean and SEM.

### Damage to the barrier function of the BBB by NMO-IgG in vitro

To evaluate whether NMO-IgG affects the barrier function of the BBB on the vascular side or the brain parenchymal side, we constructed the static *in vitro* BBB model which allowed long-term measurement of real-time TEER by means of an automated cell monitoring system. After addition of NMO-IgG or Control IgG with either the vascular side, the brain parenchymal side, or both sides, the TEER values were measured by using an automated cell monitoring device that recorded the TEER value every minute for 5 consecutive days. Within 24 hours of application of NMO-IgG, TEER values in all groups had begun to decrease compared to Control IgG, and at 72 hours they had significantly decreased in all groups (Fig. 4, A and B). Application of NMO-IgG to both the vascular and brain parenchymal sides resulted in the lowest TEER values of all groups. TEER values with brain parenchymal application of NMO-IgG were significantly lower than TEER values with vascular application of NMO-IgG.

**Figure 4.**
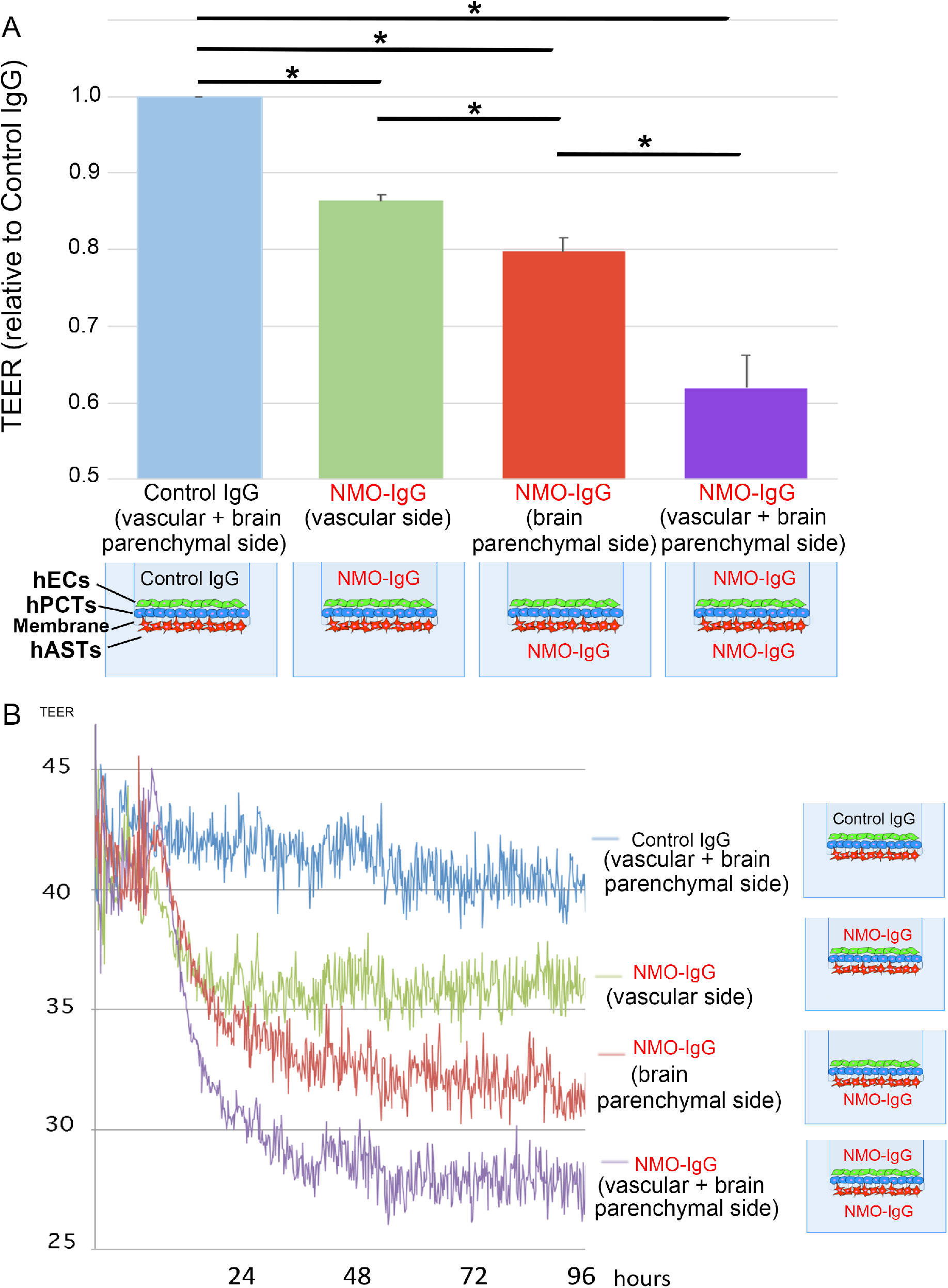
Damage to the barrier function of the BBB by NMO-IgG in *vitro*. **(6)** Application of NMO-IgG to either the vascular side or the brain parenchymal side or both sides of the static BBB model significantly decreased the TEER value relative to that of Control IgG at 72 hours. \**P* < 0.05 by unpaired t-test (*n* = 3 per group). All data are expressed as mean and SEM. **(B)** Real-time TEER measurement by cellZscope showed that the TEER values had started to decrease within 24 hours of application of NMO-IgG in all groups, and the declining trend continued for 48 hours.

### Effects of satralizumab on the barrier function of the BBB in vitro

To evaluate whether satralizumab affects the BBB dysfunction induced by NMO-IgG, we used the static *in vitro* BBB model incorporating hEC/hPCT/hAST triple co-culture with either the vascular side or the brain parenchymal side or both sides exposed to NMO-IgG. After addition of satralizumab plus NMO-IgG or Control IgG to either the vascular side or the brain parenchymal side or both sides, TEER values were measured by the automated cell monitoring device as mentioned above. TEER values at 72 hours were significantly higher under conditions where both the vascular side and the brain parenchymal side were exposed to satralizumab plus NMO-IgG, than under conditions of NMO-IgG alone (Fig. 5A). The inhibiting effect of satralizumab on barrier dysfunction was almost the same when satralizumab was applied to the brain parenchymal side as it was when applied to the vascular side (Fig. 5, B and C). Application of satralizumab to both the vascular and brain parenchymal sides had the highest inhibiting effect on barrier dysfunction among the three conditions (Fig. 5, A to C).

**Figure 5.**
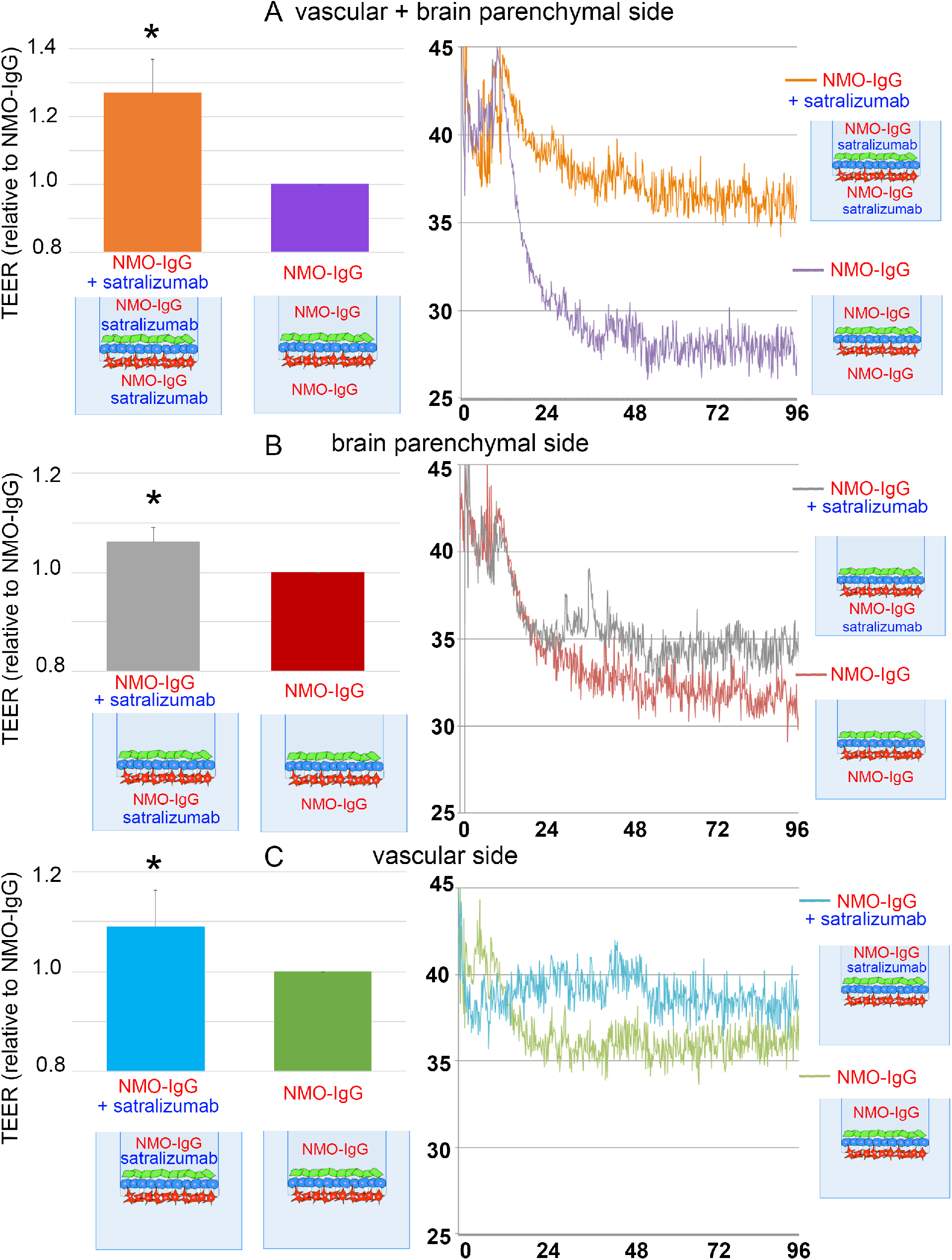
Effects of satralizumab on the barrier function of the BBB in *vitro*. (**A** to **C**, left panels) After addition of satralizumab and NMO-IgG to either the vascular side or the brain parenchymal side or both sides, the TEER values under conditions of satralizumab plus NMO-IgG were significantly higher than under conditions of NMO-IgG alone at 72 hours. \**P* < 0.05 by unpaired t-test (*n* = 3 per group). All data are expressed as mean and SEM. (**A** to **C**, right panels) Real-time TEER measurement by cellZscope showed a declining trend in TEER values under conditions of satralizumab plus NMO-IgG in all groups, but the decline remained less than that for NMO-IgG alone for 96 hours.

### Effects of IL-6 receptor blockade on the barrier function of the BBB in EAE mice

Immunohistochemical analysis on Day 15 showed that leakage of albumin and IgG into the spinal cord was higher in EAE mice than in Control mice (Fig. 6, A and B). This leakage into the CNS indicates the increased permeability of the BBB. Treatment of EAE mice with MR16-1 significantly prevented these leakages into the spinal cord. As regards the inhibiting effect on barrier dysfunction with application of satralizumab on the vascular side, we showed that sera from EAE mice mildly reduced the barrier function of endothelial cells in monoculture (Fig. S1) Serum from EAE mice at onset of EAE on Day 16 after immunization significantly decreased the TEER value of a monolayer of mouse primary BMECs. Anti-IL-6 receptor antibody significantly prevented this reduction.

**Figure 6.**
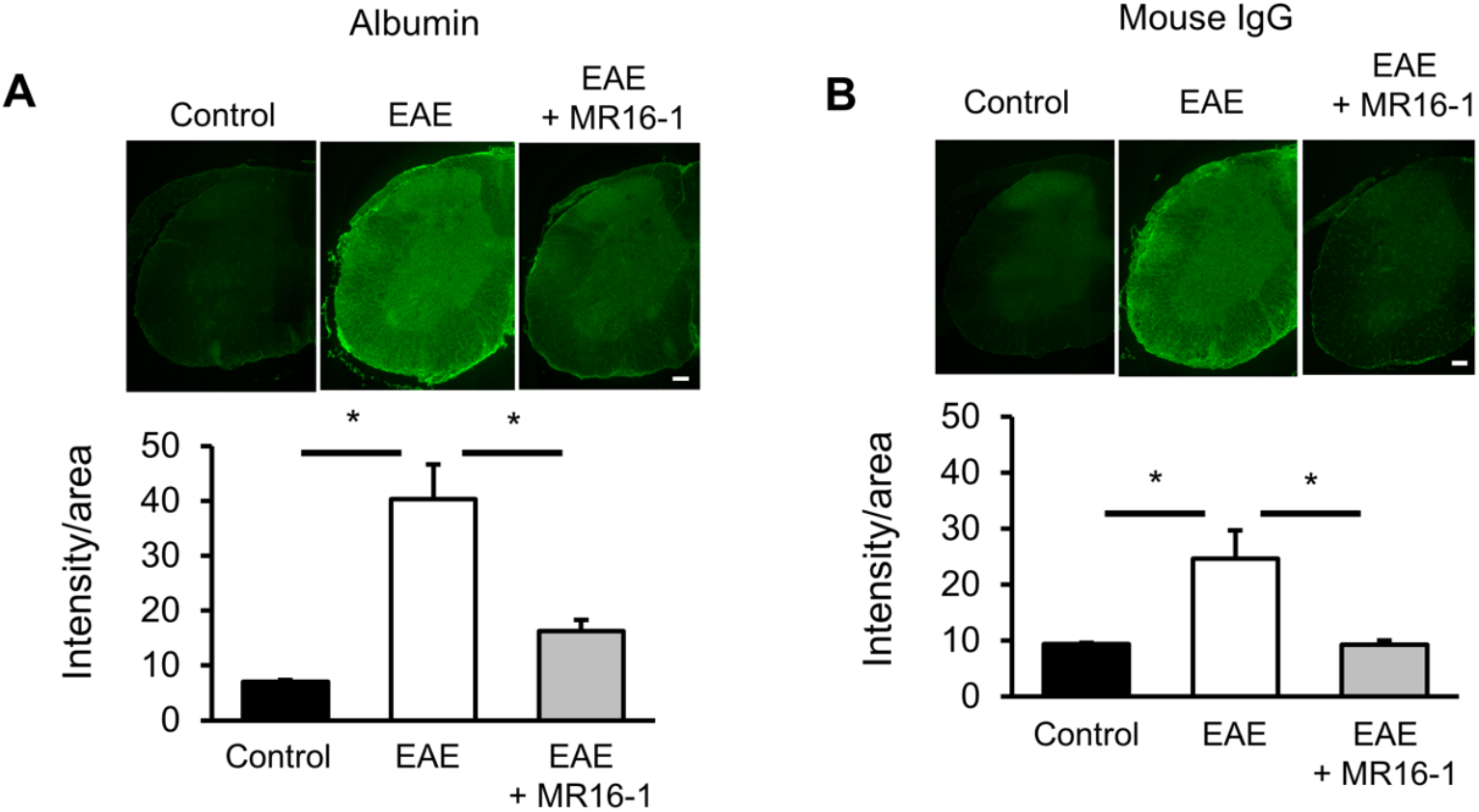
Effects of IL-6 receptor blockade on BBB permeability *in vivo*. (**A** and **B**) Representative images showing immunohistochemical staining for albumin (**A**) and IgG (**B**) in the spinal cord on Day 15 after immunization. Leakage of albumin and IgG into the spinal cord was higher in EAE mice than in Control mice, and was significantly reduced by treatment with anti-IL-6 receptor antibody (MR16-1). \**P* < 0.05 by Tukey’s multiple comparison test (*n* = 3-6 per group). Scale bars = 100 μm. All data are expressed as mean and SEM.

**Fig. S1.**
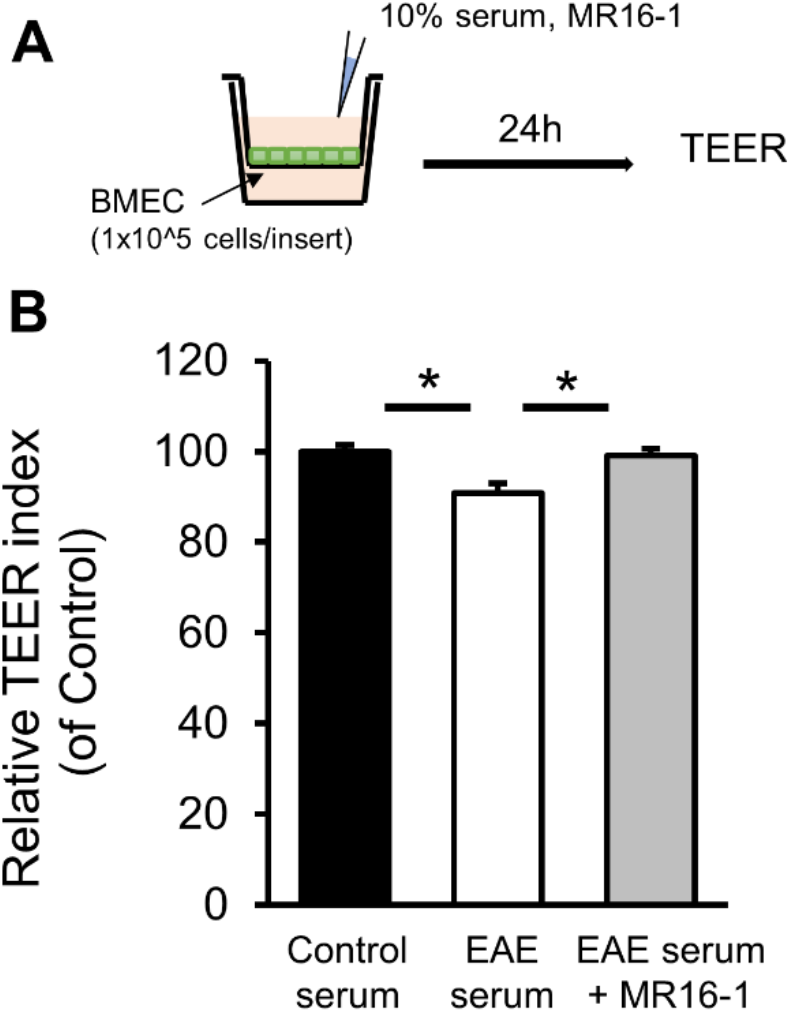
Effects of IL-6 receptor blockade on the barrier dysfunction induced by serum from EAE mice. (**A**) Graphical representation of study design used to evaluate the effect of anti-IL-6 receptor antibody on the TEER of brain microvascular endothelial cells (BMECs). (**B**) Serum from EAE mice at onset of EAE on Day 16 after immunization significantly decreased the TEER value of a monolayer of mouse primary BMECs. Anti-IL-6 receptor antibody significantly prevented this reduction.\**P* < 0.05 by Tukey’s multiple comparison test (*n* = 8 per group). All data are expressed as mean and SEM.

### Intracerebral transferability of satralizumab in the presence of NMO-IgG in vitro

To compare the microvolume translocation of satralizumab and NMO-IgG across the BBB with that of Control IgG, we constructed the *in vitro* BBB models in which measurement of microvolumes of satralizumab and IgG translocation through the BBB could be detected by Odyssey Infrared-Imaging System and ELISA. We evaluated the apparent permeability coefficients of the BBB (P_app_; mm/s) with respect to satralizumab and Control IgG in a static *in vitro* BBB model. Labeled satralizumab or IgG2 (Control IgG) was added to the hEC (vascular) side. Then the IgG that was translocated to the brain parenchymal side was detected by an infrared imaging system, and the apparent BBB permeability coefficients for satralizumab and Control IgG were calculated. The P_app_ with respect to satralizumab was almost three times the P_app_ with respect to Control IgG (Fig. 7A).

**Figure 7.**
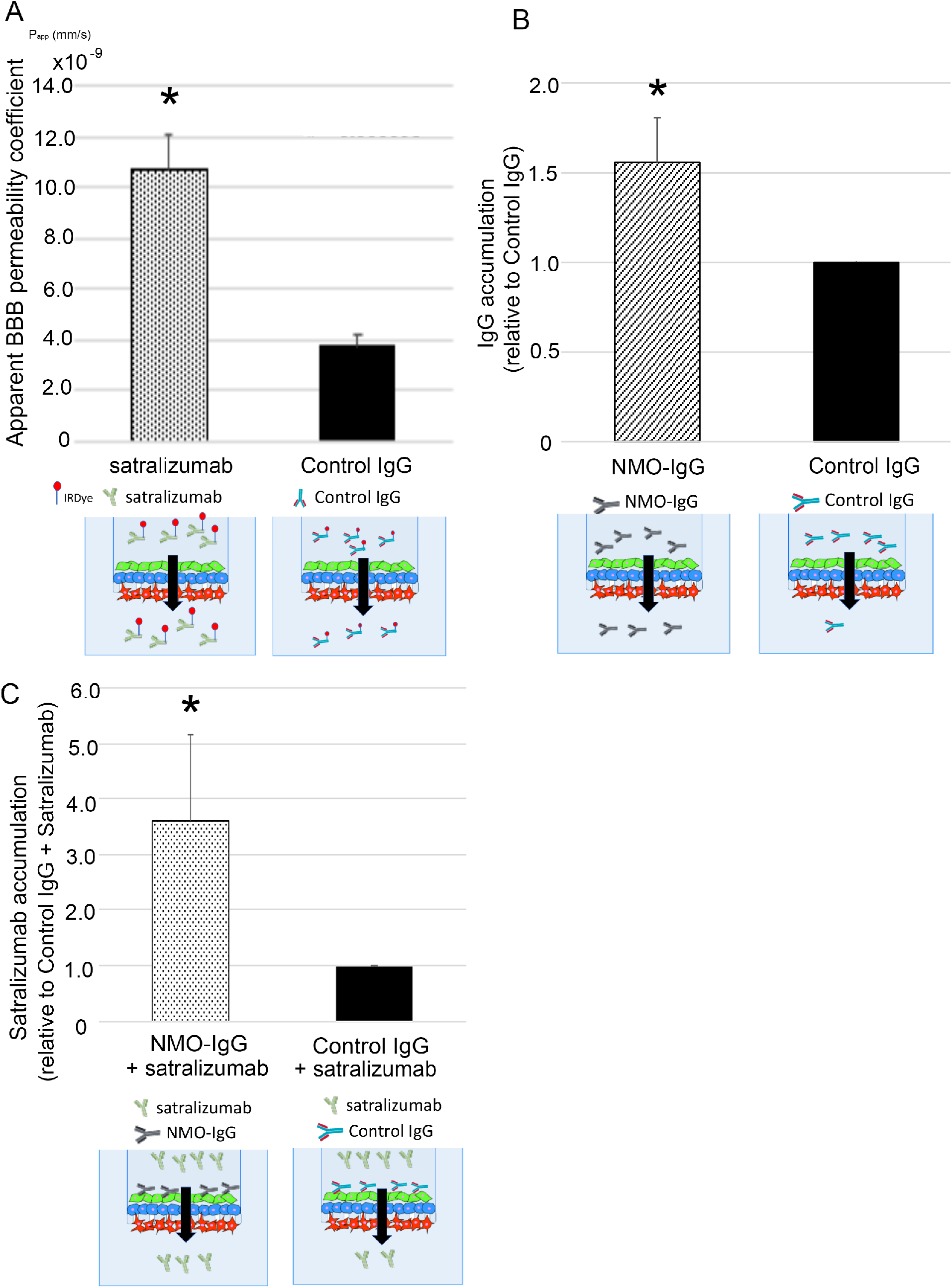
Intracerebral transferability of satralizumab in the presence of NMO-IgG *in vitro*. **(A)** Analysis of microvolume IgG translocation through the BBB by the Odyssey Infrared Imaging System revealed that the BBB P_app_ for satralizumab was almost three times that for Control IgG. \**P* < 0.05 by unpaired t-test (*n* = 6 per group). **(B)** Analysis of microvolume IgG translocation through the BBB by the spectrophotometer revealed that the relative accumulation of IgG for NMO-IgG was almost 1.5 times that for Control IgG. \**P* < 0.05 by unpaired t-test (*n* = 6 per group). **(C)** ELISA with anti-satralizumab antibody showed that the relative accumulation of satralizumab for satralizumab + NMO-IgG was significantly increased to almost three times that for satralizumab + Control IgG.\**P* < 0.05 by unpaired t-test (*n* = 8 per group). All data are expressed as mean and SEM.

After exposing the vascular side of the triple co-culture BBB model to NMO-IgG or Control IgG, the translocated IgG was detected by human IgG detection ELISA kit. The total amounts of accumulated IgG for NMO-IgG and Control IgG were calculated, normalized with respect to Control IgG, and reported as “IgG accumulation”. The intracerebral transferability of NMO-IgG was almost 1.5 times that of Control IgG (Fig. 7B).

In a previous study, we found the transfer rate of MR16-1 across the BBB in EAE mice to be almost 30 times that of normal mice (*32*). Thus, we explored whether the intracerebral transferability of satralizumab would be similarly increased in the presence of NMO-IgG. After exposing the vascular side of the triple co-culture BBB model to satralizumab plus NMO-IgG or to satralizumab plus Control IgG, the accumulated satralizumab was measured by ELISA with anti-satralizumab antibody. The total amount of satralizumab for “NMO-IgG plus satralizumab” was normalized with respect to “Control IgG plus satralizumab” and reported as “satralizumab accumulation”. The relative satralizumab accumulation for “NMO-IgG plus satralizumab” was significantly increased to almost three times that for “Control IgG plus satralizumab” (Fig. 7C).

## Discussion

Here, we successfully generated ideal *in vitro* and *in ex-vivo* BBB models for evaluating barrier function, leukocyte transmigration and intracerebral transferability of IgGs and satralizumab across the BBB utilizing the newly established triple co-culture system of temperature-sensitive conditionally immortalized human BBB cell lines (Fig.1). In *in vitro* studies, application of satralizumab to the vascular and brain parenchymal sides of the model suppressed the transmigration of total PBMCs and CD4^+^ and CD8^+^ cells, the transmigration of which was enhanced by NMO-IgG (Fig. 2). Then, we have shown that blockade of IL-6 signaling in EAE mice suppressed the migration of CD4-positive T cells into the spinal cord, prevented the increase in BBB permeability, and prevented the onset of myelitis (Fig. 3 and 6). In addition, NMO-IgG was found to cause significant barrier dysfunction, which was strongest when NMO-IgG was applied to both the vascular and brain parenchymal sides of the model; the effect was not as strong with application of NMO-IgG to the brain parenchymal side but it was stronger than with vascular application (Fig. 4). On the other hand, application of satralizumab inhibited NMO-IgG-induced barrier dysfunction; application of satralizumab to the vascular and brain parenchymal sides had the highest effect on inhibiting NMO-IgG-induced barrier dysfunction, while the effect of applying satralizumab to the brain parenchymal side was almost the same as with vascular application (Fig. 5). We also demonstrated that the intracerebral transferability of satralizumab was about three times that of Control IgG and in the presence of NMO-IgG the intracerebral transferability of satralizumab was even further increased to more than that of NMO-IgG itself (Fig. 7).

In an earlier study into the pathophysiology of NMOSD at the BBB, we found that autoantibodies to GRP78 in NMO-IgG activate NF-κB signals in vascular endothelial cells and increase BBB permeability, leading to attack by NMO-IgG on astrocytes on the CNS side of the BBB (*17*). We also showed that NMO-IgG on the CNS side induces IL-6 expression in astrocytes and that IL-6 trans-signaling affects endothelial cells and modifies the properties of the BBB, including inducing the expression of several chemokines (CCL2 and CXCL8) and decreasing the expression of claudin-5 as well as increasing the permeability with respect to solutes and increasing the transmigration of PBMCs (*11*). In the current study, we demonstrated that brain parenchymal application of NMO-IgG decreased the barrier function more strongly than did vascular application (Fig. 4), indicating that the reduction of barrier function by IL-6 signaling on the CNS side is much more than by NMO-IgG on the vascular side.

Therefore, with respect to the pathophysiology of NMOSD at the BBB, there appear to be several steps on both sides of the BBB involved in the onset of NMOSD (Fig. 8A). First, NMO-IgG (anti-GRP78 or an unknown antibody or antibodies) activates NF-κB signals in endothelial cells. Second, NMO-IgG decreases the barrier function on the vascular side. Third, NMO-IgG increases the intracerebral transferability of NMO-IgG itself. Fourth, NMO-IgG attacks the AQP4 of astrocytes and induces IL-6 expression in astrocytes. Fifth, IL-6 signaling affects endothelial cells on the CNS side. Sixth, IL-6 signaling more strongly decreases the barrier function on the CNS side than on the vascular side. Seventh, IL-6 signaling induces the expression of several chemokines (CCL2 and CXCL8) in endothelial cells. Eight, the induced chemokines enhance infiltration of inflammatory cells. Finally, NMOSD develops.

**Figure 8.**
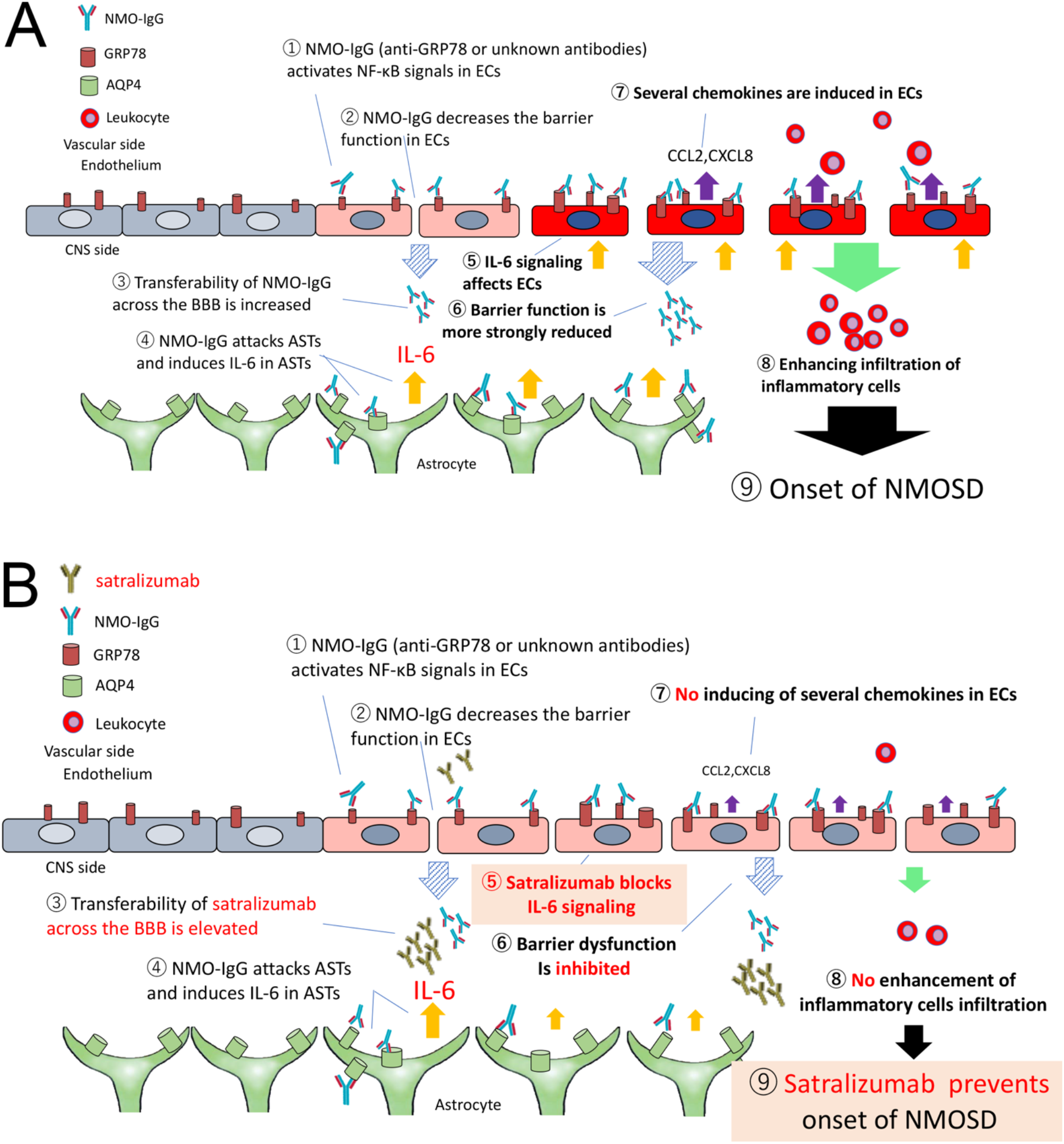
The pathophysiology of NMOSD at the BBB (A) and the action mechanism of satralizumab at the BBB (B). **(A)** There are several steps on both sides of the BBB involved in the onset of NMOSD. First, NMO-IgG (anti-GRP78 or an unknown antibody or antibodies) activates NF-κB signals in endothelial cells (ECs). Second, NMO-IgG decreases the barrier function on the vascular side. Third, NMO-IgG increases the intracerebral transferability of NMO-IgG itself. Fourth, NMO-IgG attacks the AQP4 of astrocytes (ASTs) and induces IL-6 expression in astrocytes. Fifth, IL-6 signaling affects endothelial cells on the CNS side. Sixth, IL-6 signaling more strongly decreases the barrier function on the CNS side than on the vascular side. Seventh, IL-6 signaling induces the expression of several chemokines (CCL2 and CXCL8) in endothelial cells. Eight, the induced chemokines enhance infiltration of inflammatory cells. Finally, NMOSD develops. **(B)** First, NMO-IgG (anti-GRP78 or an unknown antibody or antibodies) activates NF-κB signals in endothelial cells. Second, NMO-IgG decreases the barrier function on the vascular side. Third, NMO-IgG elevates the intracerebral transferability of satralizumab more than NMO-IgG. Fourth, NMO-IgG attacks the AQP4 of astrocytes and induces IL-6 expression in astrocytes. Fifth, satralizumab blocks IL-6 signaling on the CNS side. Sixth, satralizumab inhibits the reduction in barrier function by blockade of IL-6 signaling. Seventh, blockade of IL-6 signaling by satralizumab suppresses the expression of several chemokines (CCL2 and CXCL8) in endothelial cells. Eight, satralizumab inhibits infiltration of inflammatory cells. Finally, satralizumab prevents the onset of NMOSD.

To explore the action mechanism of satralizumab at the BBB in NMOSD, we evaluated the intracerebral transferability of satralizumab. Satralizumab was found to have potential intracerebral transferability almost three times that of Control IgG (Fig. 7A) and NMO-IgG was found to have intracerebral transferability almost 1.5 times that of Control IgG (Fig. 7B). We then found that the intracerebral transferability of satralizumab in the presence of NMO-IgG was significantly increased to almost three times that in the presence of Control IgG (Fig. 7C). These results mean that the intracerebral transferability of satralizumab is enhanced by NMO-IgG and that satralizumab can pass though the BBB and affect the CNS side.

We also demonstrated that satralizumab suppressed the decrease in barrier function induced by NMO-IgG (Fig. 5) and prevented NMO-IgG-induced enhancement of inflammatory cell infiltration (Fig. 2). In addition, administration of anti-IL-6 receptor antibody to mice at 1 week after induction of EAE prevented T cell migration and the development of EAE without inhibition of T cell differentiation (Fig. 3). Blockade of IL-6 signaling suppressed the migration of CD4-positive T cells into the spinal cord, prevented the increase in BBB permeability, and prevented the onset of the myelitis in EAE mice.

Consequently, with regard to the action mechanism of satralizumab at the BBB in NMOSD, it can be said that satralizumab can pass through the BBB in NMOSD and that blockade of IL-6 from astrocytes on the CNS side suppresses the BBB dysfunction and the induction of inflammatory cell infiltrates, leading to prevention of the onset of NMOSD (Fig. 8B). To be more specific, first, NMO-IgG (anti-GRP78 or an unknown antibody or antibodies) activates NF-κB signals in endothelial cells. Second, NMO-IgG decreases the barrier function on the vascular side. Third, NMO-IgG elevates the intracerebral transferability of satralizumab more than NMO-IgG. Fourth, NMO-IgG attacks the AQP4 of astrocytes and induces IL-6 expression in astrocytes. Fifth, satralizumab blocks IL-6 signaling on the CNS side. Sixth, blockade of IL-6 signaling by satralizumab inhibits the reduction of barrier function. Seventh, blockade of IL-6 signaling by satralizumab suppresses the expression of several chemokines (CCL2 and CXCL8) in endothelial cells. Eight, satralizumab inhibits infiltration of inflammatory cells. Finally, satralizumab prevents the onset of NMOSD.

This study also indicated that the effect of satralizumab in inhibiting barrier dysfunction was almost the same with application to the brain parenchymal side as it was with application to the vascular side, although application to both the vascular and brain parenchymal sides had the highest inhibiting effect on barrier dysfunction (Fig. 5). On the other hand, our previous study (*11*) and the current study showed that the application of NMO-IgG to the vascular side did not induce leukocyte transmigration and, as a consequence, application of NMO-IgG plus satralizumab to the vascular did not show an inhibiting effect on leukocyte transmigration, although the application of NMO-IgG plus satralizumab (or IL-6 blockade) to the brain parenchymal side or to both sides strongly suppressed leukocyte transmigration. The reason for the differences in barrier dysfunction and leukocyte transmigration seen between the vascular and CNS sides is still unsolved in this study. As regards the inhibiting effect on barrier dysfunction with application of satralizumab on the vascular side, we showed that sera from EAE mice mildly reduced the barrier function of endothelial cells in monoculture (Fig. S1). This result suggested that satralizumab (or IL-6 blockade) may directly or indirectly modify the barrier effect of endothelium on the vascular side.

We also showed that NMO-IgG increased the intracerebral transferability of satralizumab and IgG at the BBB. Because we previously found that NMO-IgG increased the permeability of the BBB with respect to solutes (*11*), the transportation of satralizumab might depend on this increased permeability. Recently, there have been several studies to identify determinants of the permeability-independent transport of immunoglobulin across the BBB (receptor-based endothelial transcytosis) (*33-39*). Several candidate molecules (transferrin receptor, insulin receptor, Fc-receptor of neonates [FCRN], and LDL receptor-related protein [LRR]) have been characterized and constitute the focus of ongoing research. In our experiments it is unresolved whether NMO-IgG increased the intracerebral transferability via permeability-dependent translocation or receptor-based endothelial transcytosis.

Because the therapeutic mechanisms of satralizumab at the BBB are as well-adapted for action in the acute phase of NMOSD by suppressing leukocyte migration as they are in the recurrence prevention period by inhibiting the barrier dysfunction, treatment with satralizumab is a promising strategy both to reduce the frequency of NMOSD attacks and to treat acute damage. Given the effect of satralizumab on BBB integrity, it may be a new option in the treatment of other conditions such as autoimmune optic neuritis and encephalomyelitis such as neuro-Behcet syndrome, CNS lupus, anti-NMDA receptor encephalitis, and Vogt-Koyanagi-Harada disease (*40–42*), all of which induce BBB breakdown and increase IL-6 concentration in the CSF.

## Materials and Methods

### Study design

This was an experimental laboratory study designed to evaluate the effect of satralizumab on BBB disruption induced by NMO-IgG. This study used human IgG which was obtained from pooled serum collected from NMOSD patients and healthy volunteers.

First, we constructed a flow-based dynamic BBB model incorporating hEC/hPCT/hAST triple co-culture, and analyzed migrating cells by flow cytometry. Then we performed *in ex-vivo* experiments to evaluate the effect of anti-IL-6 receptor antibody on leukocyte migration into the spinal cords of EAE mice. In addition to *in vivo* assays, we evaluated the effect of satralizumab on NMO-IgG-induced transmigration of leukocytes in an *in vitro* BBB model.

Second, we evaluated whether NMO-IgG affected the barrier function, especially BBB permeability, of an *in vitro* BBB model incorporating hEC/hPCT/hAST triple co-culture. We also assessed the effect of satralizumab on BBB dysfunction induced by NMO-IgG. BBB permeability was evaluated by TEER values measured by an automated cell monitoring device. Then we performed *in vivo* experiments to evaluate the effect of anti-IL-6 receptor antibody on BBB permeability in EAE mice. BBB permeability *in vivo* was assessed by immunohistochemical analysis of spinal cords of mice. Furthermore, to assess the direct effect of the anti-IL-6 receptor antibody on BBB function, we evaluated the influence of anti-IL-6 receptor antibody on the permeability of a monolayer of mouse primary brain microvascular endothelial cells stimulated by serum from EAE mice.

Finally, we measured the intracerebral transferability of satralizumab *in vitro* by using ELISA and the Odyssey Infrared Imaging System. Figure legends indicate sample sizes and statistical tests used. For all *in vivo* experiments, subjects were randomly assigned to the experimental groups before experiments.

### Human subjects

The Institutional Review Boards of Yamaguchi University Graduate School of Medicine and Chugai Pharmaceutical Co., Ltd approved all study protocols, and signed informed consent was obtained from each blood donor.

### NMO-IgG, Control IgG, and satralizumab

NMO-IgG was the IgG fraction isolated from pooled serum collected from 10 patients with NMOSD, and Control IgG came from pooled serum collected from healthy volunteers at the Yamaguchi University Graduate School of Medicine. Both NMO-IgG and Control IgG were purified by protein G affinity and adjusted for assays by extensive dialysis. Satralizumab was prepared at Chugai Pharmaceutical Co., Ltd. We used NMO-IgG, Control IgG, and satralizumab at a final concentration of 100 μg/mL.

### Cell culture

Conditionally immortalized human microvascular endothelial cells (hECs; TY10), pericytes (hPCTs), and astrocytes (hASTs) were developed by transfection with temperature-sensitive SV40 large T antigen (ts-SV40-LT). These cells proliferate at 33°C; at 37°C proliferation ceases and they differentiate into mature cells. hECs used were adult human brain microvascular endothelial cells transfected and immortalized with a plasmid expressing ts-SV40-LT as previously described (*28, 29*). hECs were grown in endothelial cell growth medium (EGM-2 Bulletkit; Lonza, Basel, Switzerland) supplemented with 10% fetal bovine serum (FBS), 100 U/mL penicillin (Sigma Aldrich, St. Louis, MO, USA), and 100 μg/mL streptomycin (Sigma Aldrich). hASTs were grown in Astrocyte Medium (ScienCell Research Laboratories, Carlsbad, CA, USA) containing 10% heat-inactivated FBS and 100 μg/mL streptomycin (Sigma Aldrich). hPCTs were maintained in Dulbecco’s-modified Eagle’s-medium (DMEM) (Gibco BRL) supplemented with 10% (v/v) heat-inactivated FBS and antibiotics (100 UI/mL penicillin G sodium, 100 μg/mL streptomycin sulfate). Astrocyte medium was used as the co-culture medium. All cells were maintained in 5% carbon dioxide at 33°C. All analyses were performed 1 or 2 days after temperature was shifted from 33°C to 37°C.

### Triple co-culture system

hPCTs and hASTs were co-cultured on Transwell insert membranes having 3 μm pores (Corning Life Sciences, Tewksbury, MA, USA), with hPCTs on the luminal side and hASTs on the abluminal side. hECs were cultured in Nunc dishes with an UpCell surface (Thermo Fisher Scientific, Waltham, MA, USA), which afford sheet-like detachment of confluent cells and extra-cellular matrix when the temperature is shifted to 20°C. The sheet of confluent hECs was detached and transferred onto the hPCTs co-cultured with hASTs on the insert. The polymers of the UpCell surface are slightly hydrophobic at 37°C but become hydrophilic at 20°C to form an aqueous film between cells and polymers resulting in sheet-like detachment of cells including their surrounding native extracellular matrix structures.

### Real-time monitoring system for TEER measurements with cellZscope

The triple co-cultured inserts were transferred to an automated cell monitoring system (cellZscope; CellSeed Inc., Tokyo, Japan). After addition of satralizumab and NMO-IgG or Control IgG to the vascular (hEC) side or the brain parenchymal (hAST) side or to both sides, the TEER values were measured using the cellZscope device which can record the TEER every minute for 120 hours.

### PBMC isolation

Peripheral blood mononuclear cells (PBMCs) were isolated from fresh heparinized blood of healthy subjects by density centrifugation with Lymphocyte Separation Medium (Mediatech, Herndon, VA, USA) as previously described (*11*), and used in transmigration assays within 2 hours of phlebotomy. For transmigration assays, PBMCs were resuspended at 10 × 10^6^ cells in 30 mL TEM buffer (RPMI 1640 without phenol red + 1% bovine serum albumin + 25 mM HEPES) and stained with Calcein AM (Thermo Fisher Scientific) prior to perfusion into the 3D flow chamber device following the manufacturer’s protocol (see below).

### Transmigration assay

Flow-based transmigration assays were performed in a 3D Bioflux flow chamber device (Fluxion Bioscience, San Diego, CA, USA) as previously described (*30*). In brief, this system comprises a 3D flow pump, 3D flow chamber, and 3D flow membranes. The pump delivers a programmable shear flow over a wide range (0.1-200 dyne/cm^2^) to up to eight flow devices. The 3D flow chamber (width, 30 mm; length, 70 mm; height, 8 mm) has three discrete reservoirs into which the 3D flow membranes fit completely. The 3D flow membrane is 8 mm in diameter and made of track-etched polycarbonate with 3 μm pores. These membranes were coated with rat-tail collagen I solution (50 μg/mL) (BD Biosciences, San Diego, CA, USA) and placed in a 12-well plate, whereupon hECs, hPCTs, and hASTs were triple co-cultured in Astrocyte Medium for 2 days at 33°C, after which time the membrane cultures were incubated for 1 day at 37°C. The membrane was gently transferred to the flow chamber, and 10 × 10^6^ PBMCs (total cells per assay) in 30 mL TEM (kept warm in a 37°C water bath) were perfused via peristaltic pump through the chamber at a final concentration of 333,000 cells/mL and at a shear stress of 1.5 dyne/cm^2^ resulting in a total assay time of 60 minutes. All chambers were set on a 37°C slide warmer. After PBMC perfusion, the chamber was flushed for 5 minutes with phosphate-buffered saline (PBS) to remove loose cells, maintaining the same shear stress as in the assay. Migrated PBMCs were recovered from the bottom chamber. Cells that attached to the abluminal side of the membrane and the bottom chamber were removed by a quick rinse with 0.5 mM EDTA. The migrated cells were enumerated by a hemocytometer, then normalized to migrated cell numbers determined by flow cytometry. After collection, cells were fixed for 10 minutes in 1% paraformaldehyde at room temperature, washed in PBS + 0.1 mM EDTA, followed by blocking in mouse IgG. Cells were labeled with anti-human CD45 efluor450, CD8a APC-efluor780 (eBiosciences, San Diego, CA, USA), CD3Alexa Fluor 647 (BioLegend, San Diego, CA, USA), CD19 BV711, and CD4 PE-CF594 (BD Biosciences). Data were acquired using BD FACSCanto II (BD Biosciences), and analyzed by FlowJo software (v.10.4.1; Treestar, Ashland, OR, USA).

### Measurement of microvolume IgG translocation by Odyssey Infrared Imaging System

Control IgG and satralizumab were labeled with IRDye 800CW protein (IRDye 800CW Protein Labeling Kit; LI-COR, Lincoln, NE, USA) following the manufacturer’s protocol. After exposing hECs in triple co-cultured inserts to labeled Control IgG or satralizumab, the microvolumes that translocated to the lower chamber were detected by an Odyssey Infrared Imaging System (LI-COR), and the apparent BBB permeability coefficient (P_app_; mm/s) was calculated from the degree of IgG translocation by using the following formula:

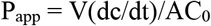

where dc/dt = flux of IgG across the membrane; V (cm^3^) = volume on the receiving side; A (cm^2^) = surface area of insert; and C_0_ (mM) = initial concentration in the donor compartment.

### Measurement of microvolume IgG translocation by spectrophotometer

NMO-IgG or Control IgG were added to hECs in triple co-cultured inserts and incubated for 12 hours. Following the manual of the Easy-Titer Antibody Assay Kit (Thermo), anti-human IgG coated beads were added to the lower well and incubated with the sample for 60 minutes. Accumulated IgG in the lower well was measured by spectrophotometer, following the manufacturer’s instructions. The total amount of accumulated IgG was normalized to 1 using Control IgG and reported as “IgG accumulation”.

### Measurement of microvolume satralizumab translocation by ELISA

After exposing hECs in triple co-cultured inserts to satralizumab plus NMO-IgG or satralizumab plus Control IgG for 24 hours, the concentration of satralizumab in the lower chamber was measured by using ELISA with anti-satralizumab antibody. The samples were run in according to the manufacture’s protocol.

Total amount of accumulated satralizumab was normalized to 1 using “Control IgG + satralizumab” and reported as “Satralizumab accumulation”.

### Animals

Female C57BL/6J mice (7 weeks old; Charles River Laboratories Japan, Inc., Kanagawa, Japan) were used. All mice were fed ordinary laboratory chow and allowed free access to water under a constant light and dark cycle of 12 hours. All animal procedures were conducted in accordance with the Guidelines for the Care and Use of Laboratory Animals at Chugai Pharmaceutical Co., Ltd, and all experimental protocols were approved by the Animal Care Committee of the institution (approval No. 18-144, 19-191) and conformed to the *Guide for the Care and Use of Laboratory Animals* published by the US National Institutes of Health.

### Experimental design of EAE mice

Experimental autoimmune encephalomyelitis (EAE) was induced in mice by subcutaneous immunization (on Day 0) with 50 μg of the myelin oligodendrocyte glycoprotein 35-55 peptide (MOG35-55; Peptide International, Louisville, KY, USA) emulsified in complete Freund’s adjuvant (Difco Laboratories, Detroit, MI, USA) supplemented with Mycobacterium tuberculosis extract H37Ra (Difco Laboratories). In addition, mice received 250 ng pertussis toxin (List Biological Laboratories, Campbell, CA, USA) intravenously on Day 0 and intraperitoneally on Day 2. Control mice were treated with complete Freund’s adjuvant and saline alone. Anti-IL-6 receptor antibody (MR16-1) was prepared using a hybridoma established in Chugai Pharmaceutical’s laboratories (*43*). The EAE mice were intraperitoneally administered MR16-1 (8 mg/mouse) on Day 7 after MOG35-55 immunization. On Day 15 or 16 after MOG35-55 immunization, spinal cords, spleens, and sera were harvested for immunohistochemistry, flow cytometry, and TEER studies.

### Clinical score assessment

EAE mice were sequentially scored for clinical signs of EAE according to the following scale: 0, no apparent disease; 1, limp tail; 2, hind limb weakness; 3, hind limb paresis; 4, hind limb paralysis; 5, hind limb and fore limb paralysis; 6, moribundity and death.

### Immunohistochemistry

Mice were anaesthetized with isoflurane, and transcardial perfusion was carried out with 20 mL of cold PBS. The L3-L5 segment of the lumbar spinal cord was removed, fixed in 4% paraformaldehyde, and placed in a 30% sucrose solution overnight. Samples were embedded in optimal cutting temperature (OCT) compound, and frozen slices of spinal cord (10 μm thick) were obtained with a cryostat. Spinal cord slices were stained by using the following primary antibodies: goat anti-albumin antibody (1:200, A90-134A; Bethyl Laboratories, Inc., Montgomery, TX, USA), biotin-conjugated donkey anti-mouse IgG antibody (1:200, 715-066-151; Jackson ImmunoResearch, West Grove, PA, USA), and rat anti-CD4 antibody (1:100, 550280; BD Pharmingen Inc., San Diego, CA, USA). After overnight incubation with primary antibodies at 4°C, spinal cord sections were incubated with secondary antibody Alexa Fluor 488-conjugated donkey anti-goat IgG (1:200, 705-546-147; Jackson ImmunoResearch) and Alexa Fluor 488-conjugated streptavidin (2 μg/mL, 016-540-084; Jackson ImmunoResearch). For CD4 staining, biotin-conjugated donkey anti-rat IgG (1:200, 712-066-153; Jackson ImmunoResearch) and Alexa Fluor 488-conjugated streptavidin (2 μg/mL, 016-540-084; Jackson ImmunoResearch) were used. Slides were mounted using Vectashield Antifade Mounting Medium (H-1200; Vector Laboratories, Burlingame, CA, USA). Spinal cord slices were randomly selected from each mouse and observed under a BZ-9000 Fluorescence Microscope (Keyence, Osaka, Japan). Positive-staining areas were calculated using BZ-II analyzer (Keyence) and CD4-positive T cells were counted with imaging analysis software (WinROOF Version 6.3.1; Mitani Corporation, Fukui, Japan)

### Flow cytometry

Spleens collected from Control mice and EAE mice were homogenized and passed through a 100 and 40 μm cell strainer to isolate mononuclear cells. Red blood cells were hemolyzed with ACK lysing buffer (Gibco, Carlsbad, CA, USA). Mononuclear cells were incubated with Mouse BD Fc Block (BD Pharmingen Inc.) before staining. For intracellular cytokine staining, mononuclear cells were stimulated for 4 hours in RPMI 1640 (Sigma, St. Louis, MO, USA) containing 10% FBS (Gibco), 55 μM 2-mercaptoethanol (Gibco), 10 mM HEPES (Sigma), 1 mM sodium pyruvate (Wako, Osaka, Japan), and 100 U /mL penicillin-streptomycin (Gibco) with 50 ng/mL phorbol 12-myristate 13-acetate (Sigma) and 1 μM ionomycin (Sigma) in the presence of 0.1% BD GolgiPlug (BD Biosciences). The cells were initially stained with FITC-conjugated anti-CD4 antibody (100510; BioLegend), and then intracellularly stained using PE-conjugated anti-IL-17A (506904; BioLegend), and APC-conjugated anti-IFN-γ (505810; BioLegend) antibodies; staining was performed with the Fixation/Permeabilization Solution Kit with BD GolgiPlug (BD Biosciences) according to the manufacturer’s protocol. For analysis of Treg cells, Fc-blocked cells were initially stained with FITC-conjugated anti-CD4 and BV421-conjugated anti-CD25 (102034; BioLegend) antibodies, and then intracellularly stained using the APC-conjugated anti-Foxp3 antibody (17-5773-80B; Invitrogen, Carlsbad, CA, USA); staining was performed with a Foxp3 Transcription Factor Staining Buffer Set (Invitrogen) according to the manufacturer’s protocol. Data were acquired using BD FACSCanto II (BD Biosciences) and analyzed using FlowJo 10.4.1 (Treestar).

### Statistical analysis

All data are expressed as mean and SEM. The statistical significance of differences was determined by using unpaired t-test, Tukey’s multiple comparison test with ANOVA, or two-way ANOVA for comparison of time course data. Probability values of less than 0.05 were considered significant. Statistical analyses were performed using IBM SPSS Statistics (International Business Machines Corporation, Armonk, NY, USA) or JMP version 11.2.1 software (SAS Institute, Cary, NC, USA).

## Supplementary Materials

Supplementary materials and methods

Fig. S1. Effects of IL-6 receptor blockade on the barrier dysfunction induced by serum from EAE mice.

## Acknowledgments

The authors thank Yoshihiro Matsumoto and Kenji Yogo of Chugai Pharmaceutical Co., Ltd for ensuring the integrity of the *in vivo* study.

## Author contributions

T.K. and R.M.R. conceptualized and designed the study. Y.T., S.F., and K.S. were involved in drafting the article. Y.T. performed real-time monitoring of TEER measurements with cellZscope. S.F. isolated PBMCs and conducted transmigration assays. Y.T. and K.M. constructed the triple co-culture system. S.F. and M.F. measured microvolume IgG and satralizumab. J.N., F.S., and Y.S. cultured the hECs, hPCTs, and hASTs and controlled the quality of these cell lines. K.S., H.T.-S., and S.M. were involved in data acquisition in EAE experiments.

## Data and material availability

All data associated with this study are present in the paper. hECs, hPCTs, and hASTs are available from T.K. under a material transfer agreement with the Yamaguchi University. MR16-1 is available under a material transfer agreement with Chugai Pharmaceutical Co., Ltd.

## Supplementary Materials

### Materials and Methods

#### Transendothelial electrical resistance studies using mouse serum

C57BL/6 Mouse Primary Brain Microvascular Endothelial Cells (BMECs, C57-6023; Cell Biologics Inc., Chicago, IL, USA) were seeded on the upper (luminal) surface of culture well insert membranes. On the following day, cells were incubated with medium containing 10% serum from Control mice or EAE mice (clinical score ≥1). MR16-1 (100 μg/mL) and soluble IL-6 receptor (100 ng/mL) were also added. After 1 day of incubation, TEER values were measured with Endohm-6 and EVOM2 (World Precision Instruments, Sarasota, FL, USA).

